# Site-specific analysis of the SARS-CoV-2 glycan shield

**DOI:** 10.1101/2020.03.26.010322

**Authors:** Yasunori Watanabe, Joel D. Allen, Daniel Wrapp, Jason S. McLellan, Max Crispin

## Abstract

The emergence of the betacoronavirus, SARS-CoV-2 that causes COVID-19, represents a significant threat to global human health. Vaccine development is focused on the principal target of the humoral immune response, the spike (S) glycoprotein, that mediates cell entry and membrane fusion. SARS-CoV-2 S gene encodes 22 N-linked glycan sequons per protomer, which likely play a role in immune evasion and occluding immunogenic protein epitopes. Here, using a site-specific mass spectrometric approach, we reveal the glycan structures on a recombinant SARS-CoV-2 S immunogen. This analysis enables mapping of the glycan-processing states across the trimeric viral spike. We show how SARS-CoV-2 S glycans differ from typical host glycan processing, which may have implications in viral pathobiology and vaccine design.

## Introduction

Severe acute respiratory syndrome coronavirus-2 (SARS-CoV-2), the causative pathogen of COVID-19^1,2^, induces fever, severe respiratory illness and pneumonia. SARS-CoV-2 utilizes an extensively glycosylated spike (S) protein that protrudes from the viral surface to bind to angiotensin-converting enzyme 2 (ACE2), the host cell receptor, to mediate cell entry^3^. The S protein is a trimeric class I fusion protein that is composed of two functional subunits responsible for receptor binding (S1 subunit) and membrane fusion (S2 subunit). Remarkably, the surface of the virally encoded envelope spike is dominated by an array of host-derived glycans with each trimer displaying 66 N-linked glycosylation sites. This extensive glycosylation has important implications for vaccine design.

As obligate parasites, many viruses exploit host-cell machinery to glycosylate their own proteins. Numerous viral envelope proteins, including HIV-1 envelope (Env), influenza hemagglutinin (HA) and Lassa virus glycoprotein complex (GPC), possess genetically encoded N-linked glycan sequons (N-X-S/T motifs, where X is any amino acid except proline). Viral glycosylation has wide-ranging roles in viral pathobiology, including mediating protein folding and stability, and shaping viral tropism. The genetically encoded sequons can be under significant selective pressure as a mechanism for immune evasion by shielding specific epitopes from antibody neutralization. However, we note the currently reported low mutation rate of SARS-CoV-2, and as yet that there have been no observed mutations to N-linked glycosylation sites^4^. Surfaces with an unusually high density of glycans can also enable immune recognition^5–7^. The role of glycosylation in immune evasion by camouflaging immunogenic protein epitopes has been well studied for other coronaviruses^4,8,9^.

As the principal antigen presented on the surface of SARS-CoV-2 virions, the S protein is a key target in vaccine design efforts. It is apparent that the viral spike will be targeted by the full assortment of vaccine strategies from nucleic-acid based approaches^10^, whereby the viral protein is expressed *in vivo*, to recombinant strategies whereby viral glycoproteins are delivered with appropriate adjuvants. Such strategies aim to elicit neutralizing adaptive immunity with an emphasis on achieving an antibody response at the sites of viral entry.

Understanding the glycosylation of recombinant viral spikes can both reveal fundamental features of viral biology and can guide vaccine design strategies and manufacturing. As glycans are enzymatically elaborated in the Golgi apparatus, some features of processed, so-called complex-type, glycosylation will necessarily be influenced by the producer cell-line. However, the presence of glycans typical of the early stages of the secretory pathway on otherwise mature glycoproteins often closely relate to fundamental features of viral spike architecture. High viral glycan density, or extensive packing of the glycan to the protein surface, can impair the normal glycan maturation pathway by steric interference of host enzymes. Under-processed oligomannose- and hybrid-type glycans are found on the majority of viral envelope proteins such as HIV-1 Env, influenza HA and Lassa virus GPC^5^. These viral glycoproteins traffic through the secretory system and the glycosylation processing of recombinant material often closely captures the glycan maturation state of the virion. This can be particularly important for viruses such as HIV-1, where viral glycans can also be targeted by neutralizing antibodies. Coronaviruses have been reported to form virions by budding into the lumen of endoplasmic reticulum-Golgi intermediate compartments (ERGIC)^11–14^. However observations of hybrid- and complex-type glycans on virally derived material suggests that the viral glycoproteins are subjected to Golgi resident processing enzymes^8,15^.

As impaired glycan maturation can be a sensitive reporter of local viral protein architecture^16^, detailed site-specific analysis has emerged as an indicator of native-like architecture and is increasingly used to compare different immunogens and in the monitoring of manufacturing processes. Importantly, in addition to these structural insights, the presence of oligomannose-type glycans on viral spike-based immunogens has also been shown to enhance trafficking of glycoprotein to germinal centers via interaction with lectins such as mannose-binding lectin^17^. It is therefore of considerable importance to understand the glycosylation of recombinant mimetics of the virus spike.

Here, we apply mass spectrometry to understand both the site-specific N-linked glycan composition and the degree of sequon occupancy on a soluble, native-like SARS-CoV-2 S protein. The native-like folding of trimeric recombinant material has been recently revealed by detailed structural analysis by cryo-electron microscopy^18,19^. We have previously validated our glycopeptide analysis methodology and applied this approach to the study of a range of other viral glycoprotein immunogens^4,20–23^, which enables cross-viral comparisons of glycosylation to be made. We report here the site-specific glycosylation at each of the 22 N-linked glycan sites found on the SARS-CoV-2 S protomer. As observed on other viral glycoproteins, there is an elevation in oligomannose- and hybrid-type glycans compared to host-derived glycoproteins, although compared to many other viruses there are still a large population of complex-type glycans displayed across the trimer surface. We also report that for each of the 22 glycan sites the occupancy is nearly fully complete, which means that any epitopes shielded from the immune system on the virus will also likely be shielded on the immunogen. Site-specific glycan analysis of SARS-CoV-2 immunogens will help guide vaccine design and manufacturing.

## Results and discussion

### Localized impairment to SARS-CoV-2 S glycan maturation

To resolve the site-specific glycosylation of SARS-CoV-2 S protein and visualize the distribution of glycoforms across the protein surface, we expressed and purified recombinant soluble material in an identical manner to that which was used to obtain the high-resolution cryo-electron microscopy (cryo-EM) structure, albeit without glycan processing blockade using kifunensine^18^. This soluble recombinant variant of the S protein contains all 22 glycans on the SARS-CoV-2 S protein (**Figure 1A**). Stabilization of the trimeric prefusion structure was achieved using the “2P” stabilizing mutations^24^ at residues 986 and 987 in addition to a C-terminal trimerization motif. This ensures that the quaternary structure remains intact during glycan processing, as in the case of HIV Env mimetics, this is known to influence glycosylation of certain sites^16,25^. Prior to analysis, supernatant containing the recombinant SARS-CoV-2 was purified using a C-terminal StrepTag followed by size-exclusion chromatography to ensure only native-like trimeric SARS-CoV-2 S protein is analyzed (**Figure 1B**). The trimeric conformation of the purified material was validated using negative stain electron microscopy (**Figure 1C**).

**Figure 1.**
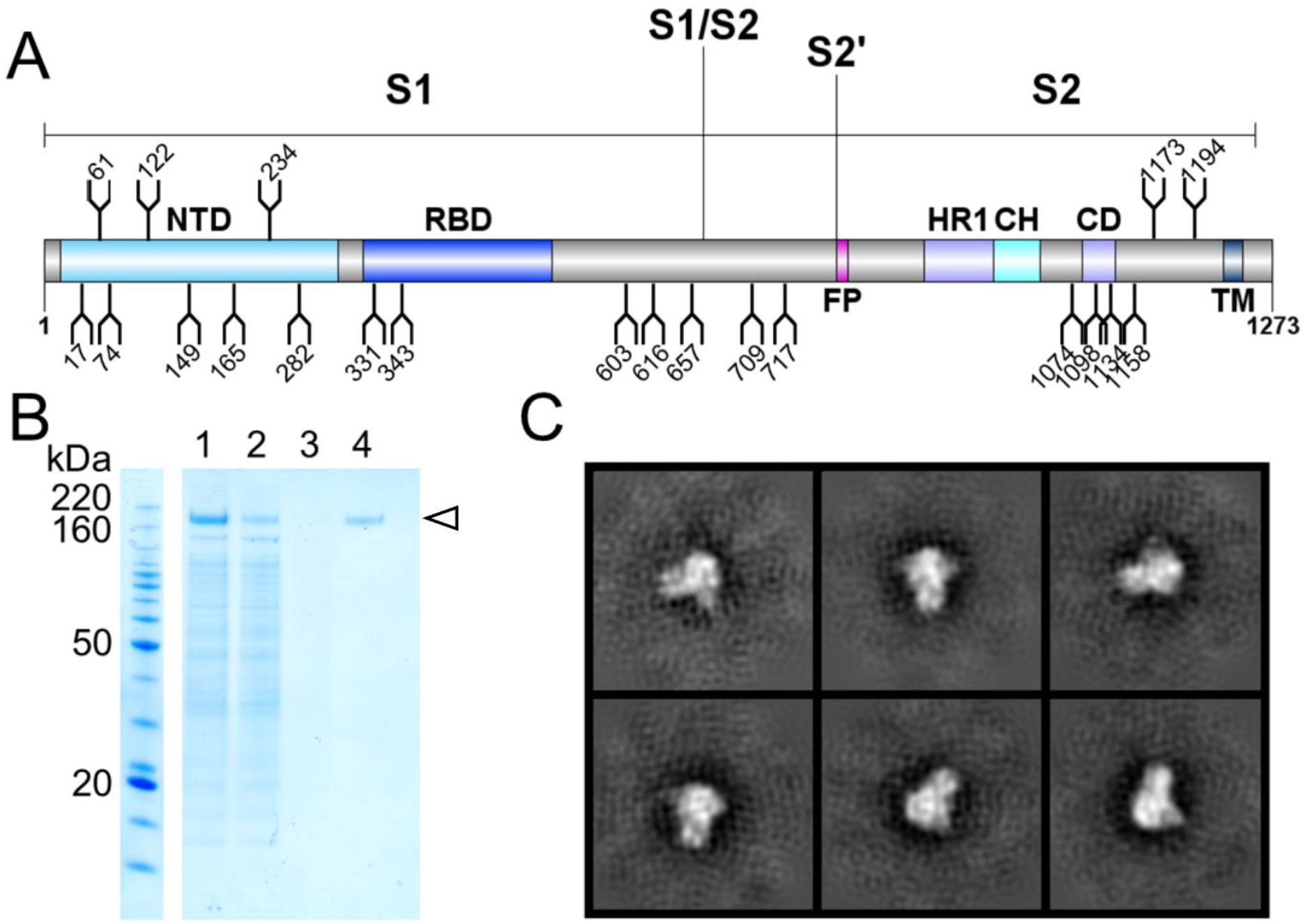
Expression and validation of SARS-CoV-2 S glycoprotein. **(A)** Schematic representation of SARS-CoV-2 S glycoprotein. The positions of N-linked glycosylation sequons (N-X-S/T, where X≠P) are shown as branches. Protein domains are illustrated: N-terminal domain (NTD), receptor-binding domain (RBD), fusion peptide (FP), heptad repeat 1 (HR1), central helix (CH), connector domain (CD), and transmembrane domain (TM). **(B)** SDS-PAGE analysis of SARS-CoV-2 S protein. Lane 1: filtered supernatant from transfected cells; lane 2: flowthrough from StrepTactin resin; lane 3: wash from StrepTactin resin; lane 4: elution from StrepTactin resin. **(C)** Negative-stain EM 2D class averages of the SARS-CoV-2 S protein. 2D class averages of the SARS-CoV-2 S protein are shown, confirming that the protein adopts the trimeric prefusion conformation matching the material used to determine the structure^18^.

Trypsin, chymotrypsin, and alpha-lytic protease were employed to generate three glycopeptide samples. These proteases cleave at different sites and were selected in order to generate glycopeptides that contain a single N-linked glycan sequon. The glycopeptide pools were analyzed by LC-MS, with high-energy collision-induced dissociation (HCD) fragmentation, and the glycan compositions at each site were determined for all 22 N-linked glycan sites on the SARS-CoV-2 S protein (**Figure 2**). A diverse range of glycan compositions were observed across the different glycosylation sites. In order to convey the main processing features at each site, the abundances of each glycan are summed into oligomannose-, hybrid- and complex-type glycosylation. In addition, the diverse signals arising from heterogeneous complex-type glycosylation are simplified by the summation of glycan intensities into a more limited range of structural categories.

**Figure 2:**
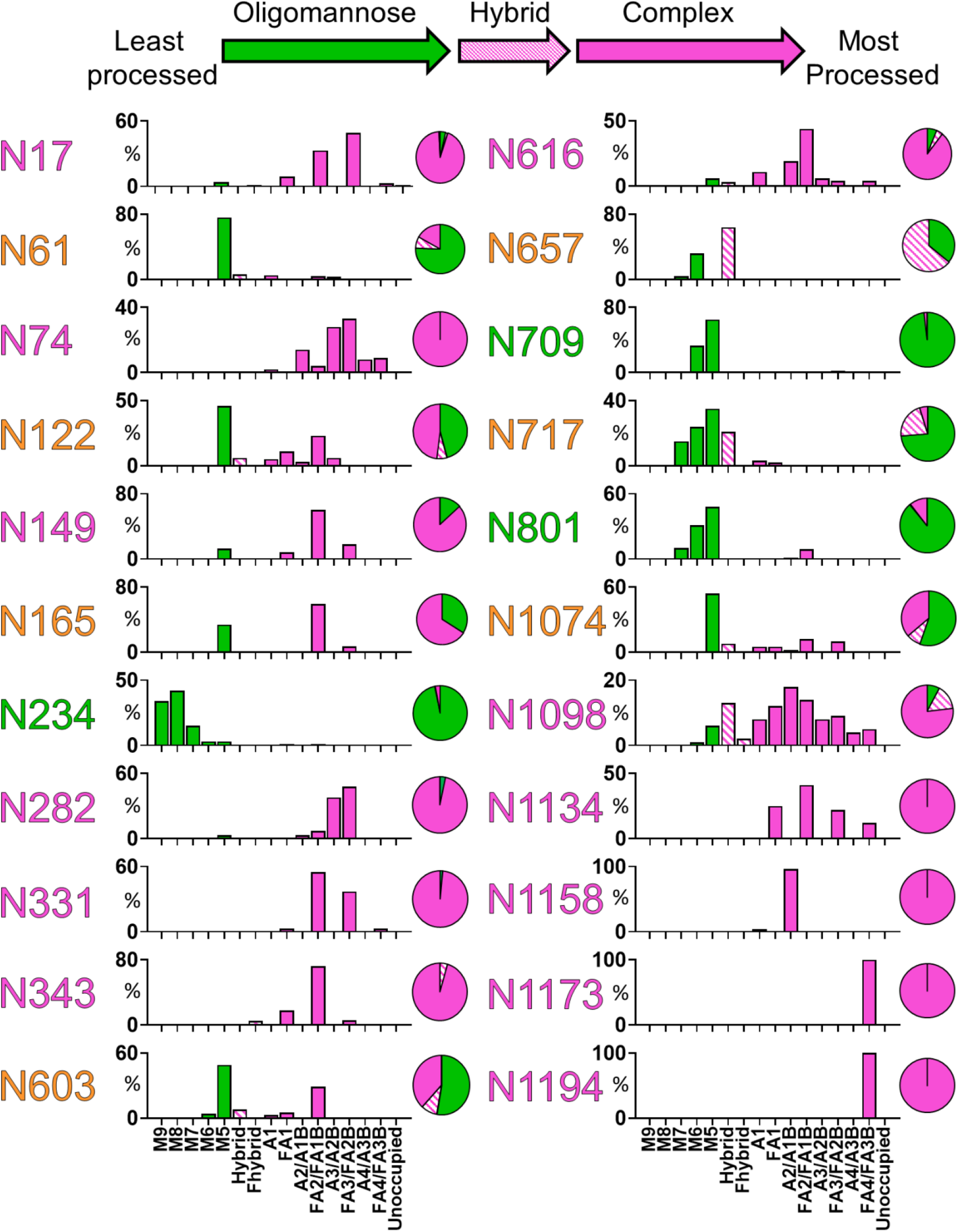
Site-specific N-linked glycosylation of SARS-CoV-2 S glycoprotein. The schematic illustrates the color code for the principal glycan types that can arise along the maturation pathway from oligomannose-, hybrid-to complex-type glycans. The graphs summarize quantitative mass spectrometric analysis of the glycan population present at individual N-linked glycosylation sites. The bar graphs represent the quantities of each glycan group with oligomannose-type glycan series (M9 to M5; Man_9_GlcNAc_2_ to Man_5_GlcNAc_2_) (green), afucosylated and fucosylated hybrid glycans (Hybrid & F Hybrid) (dashed pink), and complex glycans grouped according to the number of antennae and presence of core fucosylation (A1 to FA4) (pink). Left to right; least processed to most processed. The pie charts summarize the quantification of these glycans. Glycan sites are colored according to oligomannose-type glycan content with the glycan sites labelled in green (80−100%), orange (30−79%) and pink (0−29%).

There are three sites on SARS-CoV-2 that are predominantly oligomannose-type: N234, N709 and N801. The predominant structure observed at each site, with the exception of N234, is Man5GlcNAc2, which demonstrates that these sites are largely accessible to a α1,2-mannosidases but are poor substrates for GlcNAcT-I, which is the gateway enzyme in the formation of hybrid- and complex-type glycans in the Golgi apparatus. The stage at which processing is impeded is a signature related to the density and presentation of glycans on the viral spike. For example, the more densely glycosylated spikes of HIV-1 Env and Lassa virus GPC give rise to numerous sites dominated by Man_9_GlcNAc_2_^20–22,26^.

Interestingly, there are several sites which possess significant populations of hybrid-type glycans, most notably at N657. This phenomenon has been observed on other viral glycoproteins, such as HIV-1 Env^20,26^, and these structures are not particularly prevalent on mammalian glycoproteins^27,28^. Such hybrid-type glycans have been shown to be targeted by anti-HIV antibodies^29^ and could also be important for immunogen trafficking since they have mannose-terminating moieties^17^.

A mixture of oligomannose-type glycans and complex-type glycans can be found at sites N61, N122, N165, N603, N657, N717 and N1074 (**Figure 2**). Of the 22 sites on the S protein, 10 contain significant populations of oligomannose-type glycans, highlighting how the processing of the SARS-CoV-2 S glycans is divergent from host glycoproteins. The remaining 12 sites are dominated by processed, complex-type glycans. The predominant category of complex-type glycans observed on the S protein are fucosylated biantennary structures, very similar to those abundant on mammalian glycoproteins.

A further feature that can be interrogated using this methodology is the extent of unoccupancy of glycosylation sites, where a sequon is present but a glycan has not been attached. In HIV immunogen research, the holes generated by unoccupied glycan sites have been shown to be immunogenic and potentially give rise to distracting epitopes^30^. While unoccupied glycosylation sites were detected on SARS-CoV-2, when quantified they were revealed to form a very minor component of the total peptide pool. The efficiency of glycan site occupancy of the recombinant SARS-CoV-2 immunogen versus the counterpart for HIV may well arise due to the somewhat larger spacing of sites along the polypeptide chain. The high occupancy of N-linked glycan sequons of SARS-CoV-2 indicates that recombinant immunogens will not require further optimization to enhance site occupancy.

### Fully glycosylated model of the SARS-CoV-2 spike

Using the cryo-EM structure of the trimeric SARS-CoV-2 S protein (PDB ID 6VSB)^18^, we generated a model to map the glycosylation status of the coronavirus spike mimetic onto the experimentally determined 3D structure (**Figure 3**). This combined mass spectrometric and cryo-EM analysis reveals how the N-linked glycans occlude distinct regions across the surface of the SARS-CoV-2 spike. Glycans were modelled onto each site using predominant glycans observed (**Figure 2**) and colored according to the prevalence of oligomannose-type glycans.

**Figure 3.**
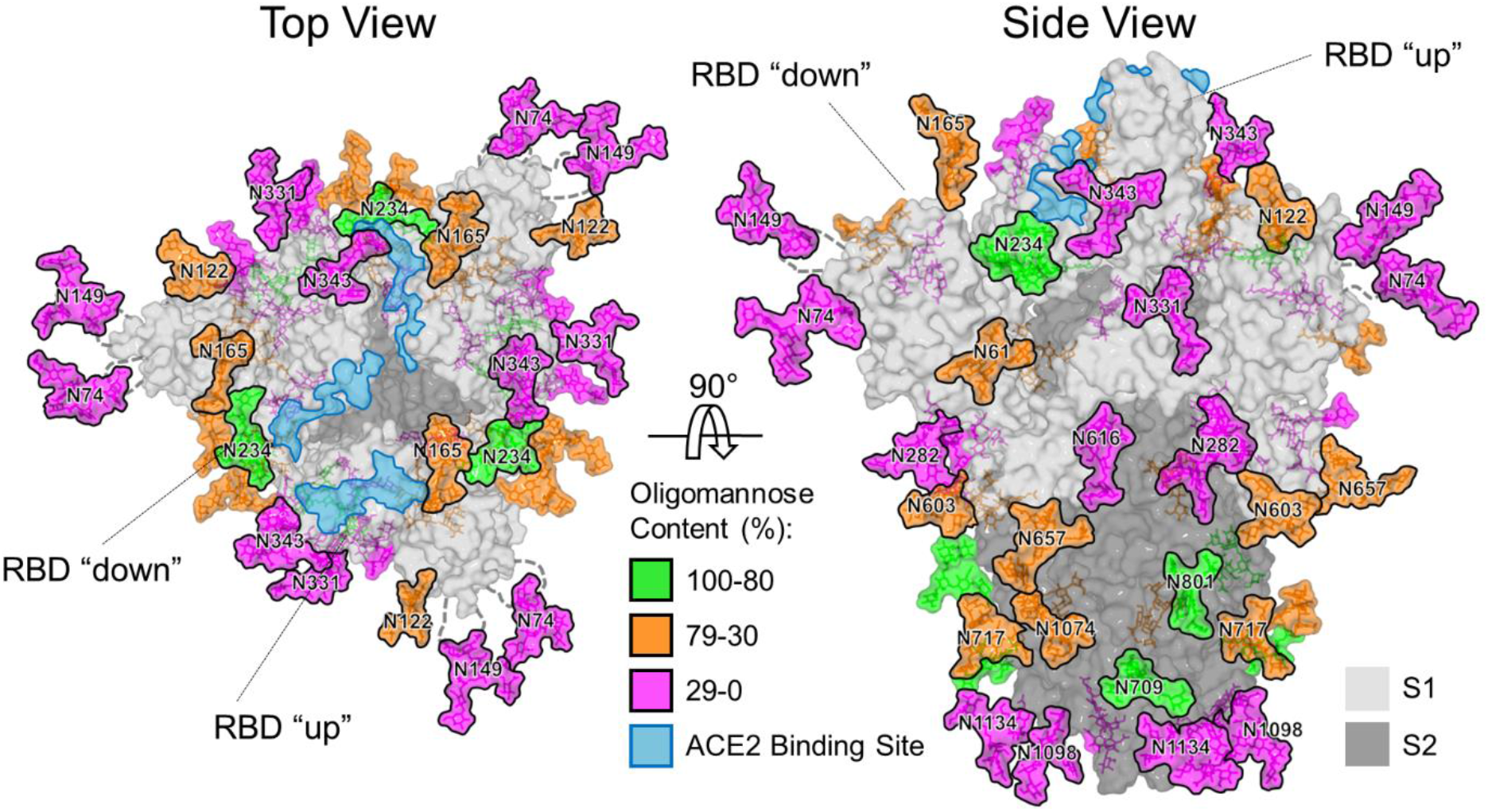
Structure-based mapping of SARS-CoV-2 S N-linked glycans. The modelling of experimentally observed glycosylation site compositions is illustrated on the prefusion structure of trimeric SARS-CoV-2 S glycoprotein (PDB ID 6VSB)^18^, with one RBD in the “up” conformation and the other two RBDs in the “down” conformation. The glycans are colored according to oligomannose content as defined by the key. ACE2 receptor binding sites are highlighted in light blue. The S1 and S2 subunits are rendered with translucent surface representation, colored light and dark grey, respectively. Note that the flexible loops on which N74 and N149 glycan sites reside are represented as dashed lines with glycan sites on the loops mapped at approximate regions.

Shielding of the receptor binding sites on the SARS-CoV-2 spike by proximal glycosylation sites (N165, N234, N343) can be observed, especially when the receptor binding domain is in the “down” conformation. The shielding of receptor binding sites by glycans is a common feature of viral glycoproteins and has been observed for SARS-CoV S^4,8^, HIV-1 Env^31^, influenza HA^32,33^, and LASV GPC^21^. Given the functional constraints of receptor binding sites and the subsequent low mutation rates of these residues, it is likely that there has been selective pressure to utilize N-linked glycans as a method to camouflage one of the most conserved and potentially vulnerable areas of their respective glycoproteins^34,35^.

It is interesting to note the absence of a specific glycan cluster that is responsible for the presence of the oligomannose-type glycans but rather there is a dispersion of these glycans across both the S1 and S2 subunits. This is in significant contrast to other viral glycoproteins, for example the density of glycans clusters in HIV have even enabled structural classification of their different modes of interaction^36^. In SARS-CoV-2 the oligomannose-type structures are probably protected, to some extent, by the protein component, as exemplified by the N234 glycan which is partially sandwiched between the N-terminal and receptor-binding domains (**Figure 3**).

We characterized the N-linked glycans on extended loop structures that were not resolved in the cryo-EM maps^18^ (N74 and N149). These were determined to be complex-type glycans, consistent with the inherent flexibility of these regions and resulting accessibility of these residues to glycan processing enzymes.

In addition to the site-specific glycosylation of the SARS-CoV-2 S protein it is also important to consider overall trends in glycosylation across the glycoprotein. The averaged compositions across all 22 glycan sites reveals that the two most common type of N-glycans on the protein are Man_5_GlcNAc_2_ (M5) and fucosylated biantennary (FA2/FA1B) glycans (**Sup. Fig. 1**). Oligomannose-type glycans comprise 32% of the total glycan pool, with hybrid-type and complex-type glycans comprising 7% and 62%, respectively (**Sup. Fig. 1**). Despite the potential impact of different local protein structure on glycan processing, the overall glycosylation of SARS-CoV-2 is comparable with SARS-CoV-1 S protein and other coronavirus S proteins^4,8,9,15^. We have previously reported that a recombinant SARS-CoV-1 S mimetic also contained 32% oligomannose-type glycans showing a remarkable conservation in glycan processing across these coronaviruses. Whilst the oligomannose-type glycan content is well above that observed on typical host glycoproteins, it is significantly lower than is found on other viral glycoproteins. For example, one of the most densely glycosylated viral spike proteins is HIV-1 Env, which contains ~60% oligomannose-type glycans^20,37^. This suggests that SARS-CoV-2 S protein is less densely glycosylated and that the glycans form much less of a shield compared with other viral glycoproteins including HIV Env and LASV GPC, which may be beneficial for the elicitation of potent neutralizing antibodies.

In addition to oligomannose-type glycans, the processing of complex-type glycans is an important consideration in immunogen engineering. Across the 22 N-linked glycosylation sites, 16% of the glycans contain at least one sialic acid residue and 48% are fucosylated (**Figure 4**). These data suggest high levels of fucosylation but low levels of sialylation, considering complex and hybrid type glycans make up 69% of the total glycans of SARS-CoV-2. Understanding the distribution of glycan modifications across the viral spike illustrates the differential susceptibility to different processing enzymes while the absolute levels can be heavily influenced by the cellular expression system utilized. We have previously demonstrated for HIV-1 Env glycosylation that the processing of complex-type glycans is driven by the producer cell but that the levels of oligomannose-type glycans were largely independent of the expression system and is much more closely related to the protein structure and glycan density^38^.

**Figure 4.**
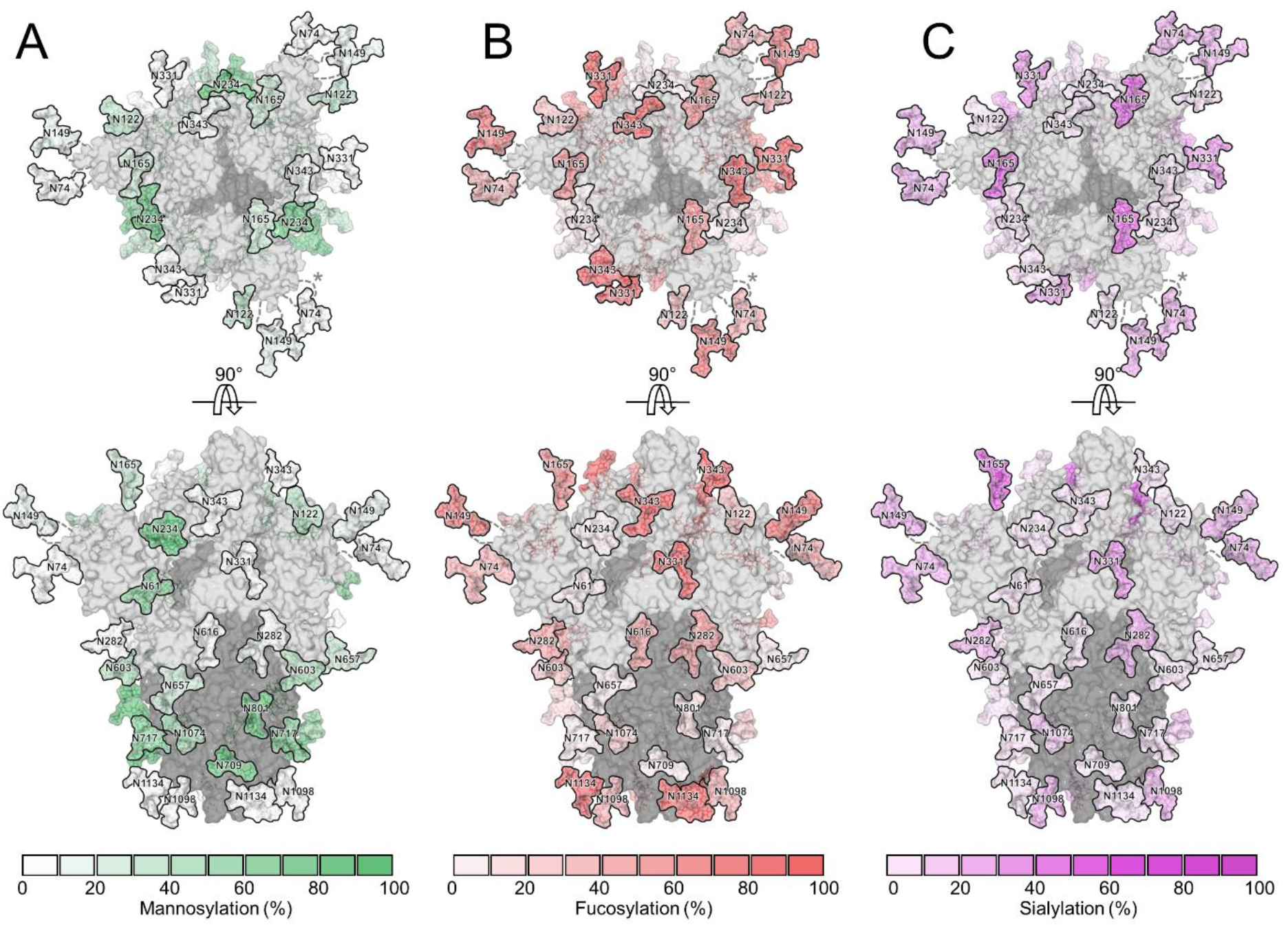
Glycosylated model of SARS-CoV-2 S glycoprotein highlighting different abundances of glycan modifications. The modelling of experimentally observed glycans, illustrated on the prefusion structure of trimeric SARS-CoV-2 S glycoprotein (PDB ID 6VSB)^18^, are highlighted according abundances of **(A)** mannosylation (Man_5-9_GlcNAc_2_), **(B)** fucosylation of the protein-proximal GlcNAc residue and **(C)** terminal sialylation. S1 and S2 subunits are colored light grey and dark grey, respectively.

## Perspectives

Our glycosylation analysis of SARS-CoV-2 offers a detailed benchmark of site-specific glycan signatures characteristic of a natively folded trimeric spike. As an increasing number of glycoprotein-based vaccine candidates are being developed, their detailed glycan analysis offers a route for comparing immunogen integrity and will also be important to monitor as manufacturing processes are scaled for clinical use. Glycosylation will therefore also be an important measure of antigen quality in the manufacture of serological testing kits, particularly as some S protein fragments may offer advantages in terms of production yield but lack effective glycan mimicry of the natively folded trimeric spike. The lower levels of mannose-terminating glycans on SARS-CoV-2 compared to many other viral spikes may indicate that glycan engineering should be considered in the scenario where first-generation glycoprotein-based vaccine candidates be poorly immunogenic. Finally, with the advent of nucleotide-based vaccines it will be important to understand how those delivery mechanisms impact immunogen processing and presentation.

## Materials and Methods

### Protein expression and purification

To express the prefusion S ectodomain, a gene encoding residues 1−1208 of SARS-CoV-2 S (GenBank: MN908947) with proline substitutions at residues 986 and 987, a “GSAS” substitution at the furin cleavage site (residues 682–685), a C-terminal T4 fibritin trimerization motif, an HRV3C protease cleavage site, a TwinStrepTag and an 8XHisTag was synthesized and cloned into the mammalian expression vector pαH. This expression vector was used to transiently transfect FreeStyle293F cells (Thermo Fisher) using polyethylenimine. Protein was purified from filtered cell supernatants using StrepTactin resin (IBA) before being subjected to additional purification by size-exclusion chromatography using a Superose 6 10/300 column (GE Healthcare) in 2 mM Tris pH 8.0, 200 mM NaCl and 0.02% NaN_3_.

### Negative-stain electron microscopy and 2D class averaging

Purified SARS-CoV-2 spike was diluted to a concentration of 0.04 mg/mL using 2 mM Tris pH 8.0, 200 mM NaCl and 0.02% NaN_3_ before being applied to a plasma cleaned CF400-Cu grid (Electron Microscopy Sciences). Protein was then stained using methylamine tungstate (Nanoprobes) before being allowed to dry at room temperature for 15 minutes. This grid was imaged in a Talos TEM (Thermo Fisher Scientific) equipped with a Ceta 16M detector. Micrographs were collected using TIA v4.14 software at a nominal magnification of 92,000×, corresponding to a calibrated pixel size of 1.63 Å/pix. CTF estimation, particle picking and 2D class averaging were performed using *cis*TEM^39^.

### Glycopeptide analysis by mass spectrometry

Three 30 μg aliquots of SARS-CoV-2 S protein were denatured for 1h in 50 mM Tris/HCl, pH 8.0 containing 6 M of urea and 5 mM dithiothreitol (DTT). Next, the S protein were reduced and alkylated by adding 20 mM iodoacetamide (IAA) and incubated for 1h in the dark, followed by a 1h incubation with 20 mM DTT to eliminate residual IAA. The alkylated Env proteins were buffer-exchanged into 50 mM Tris/HCl, pH 8.0 using Vivaspin columns (3 kDa) and digested separately overnight using trypsin chymotrypsin or alpha lytic protease (Mass Spectrometry Grade, Promega) at a ratio of 1:30 (w/w). The next day, the peptides were dried and extracted using C18 Zip-tip (MerckMilipore). The peptides were dried again, re-suspended in 0.1% formic acid and analyzed by nanoLC-ESI MS with an Easy-nLC 1200 (Thermo Fisher Scientific) system coupled to a Fusion mass spectrometer (Thermo Fisher Scientific) using higher energy collision-induced dissociation (HCD) fragmentation. Peptides were separated using an EasySpray PepMap RSLC C18 column (75 μm × 75 cm). A trapping column (PepMap 100 C18 3μM 75μM x 2cm) was used in line with the LC prior to separation with the analytical column. The LC conditions were as follows: 275 minute linear gradient consisting of 0-32% acetonitrile in 0.1% formic acid over 240 minutes followed by 35 minutes of 80% acetonitrile in 0.1% formic acid. The flow rate was set to 200 nL/min. The spray voltage was set to 2.7 kV and the temperature of the heated capillary was set to 40 °C. The ion transfer tube temperature was set to 275 °C. The scan range was 400−1600 m/z. The HCD collision energy was set to 50%, appropriate for fragmentation of glycopeptide ions. Precursor and fragment detection were performed using an Orbitrap at a resolution MS^1^= 100,000. MS^2^= 30,000. The AGC target for MS^1^=4e^5^ and MS^2^=5e^4^ and injection time: MS^1^=50ms MS^2^=54ms.

Glycopeptide fragmentation data were extracted from the raw file using Byonic™ (Version 3.5) and Byologic™ software (Version 3.5; Protein Metrics Inc.). The glycopeptide fragmentation data were evaluated manually for each glycopeptide; the peptide was scored as true-positive when the correct b and y fragment ions were observed along with oxonium ions corresponding to the glycan identified. The MS data was searched using the Protein Metrics 305 N-glycan library. The relative amounts of each glycan at each site as well as the unoccupied proportion were determined by comparing the extracted chromatographic areas for different glycotypes with an identical peptide sequence. All charge states for a single glycopeptide were summed. The precursor mass tolerance was set at 4 ppm and 10 ppm for fragments. A 1% false discovery rate (FDR) was applied. The relative amounts of each glycan at each site as well as the unoccupied proportion were determined by comparing the extracted ion chromatographic areas for different glycopeptides with an identical peptide sequence. Glycans were categorized according to the composition detected. HexNAc(2)Hex(9−5) was classified as M9 to M5. HexNAc(3)Hex(5−6)X was classified as Hybrid with HexNAc(3)Fuc(1)X classified as Fhybrid. Complex-type glycans were classified according to the number of processed antenna and fucosylation. If all of the following compositions have a fucose they are assigned into the FA categories. HexNAc(3)Hex(3-4)X is assigned as A1, HexNAc(4)X is A2/A1B, HexNAc(5)X is A3/A2B, and HexNAc(6)X is A4/A3B. As this fragmentation method does not provide linkage information compositional isomers are group, so for example a triantennary glycan contains HexNAc 5 but so does a biantennary glycans with a bisect. Any glycan containing at least one sialic acid was counted as sialylated.

### Model construction

Structural models of N-linked glycan presentation on SARS-CoV-2 were created using electron microscopy structures (PDB ID: 6VSB) along with complex-, hybrid-, and oligomannose-type N-linked glycans (PDB ID 4BYH, 4B7I, and 2WAH). The most dominant glycoform presented at each site was modelled on to the N-linked carbohydrate attachment sites in Coot^40^.

## Acknowledgements

We thank M. Dixon and M. Gowland-Pryde for supporting our work on this project during the difficulties arising from the pandemic, and G. Ould for critical reading of the manuscript. This work was funded by the International AIDS Vaccine Initiative, Bill and Melinda Gates Foundation through the Collaboration for AIDS Discovery (OPP1084519 to M.C., and 1196345 to M.C.), the NIAID (R01-AI127521 to J.S.M) and the Scripps Consortium for HIV Vaccine Development (CHAVD) (AI144462 to M.C.).

