## Supplementary material for "Site-specific analysis of the SARS-CoV-2 glycan shield": Supplmentary Information

This document includes supplementary table 1 and supplementary table 1 legend

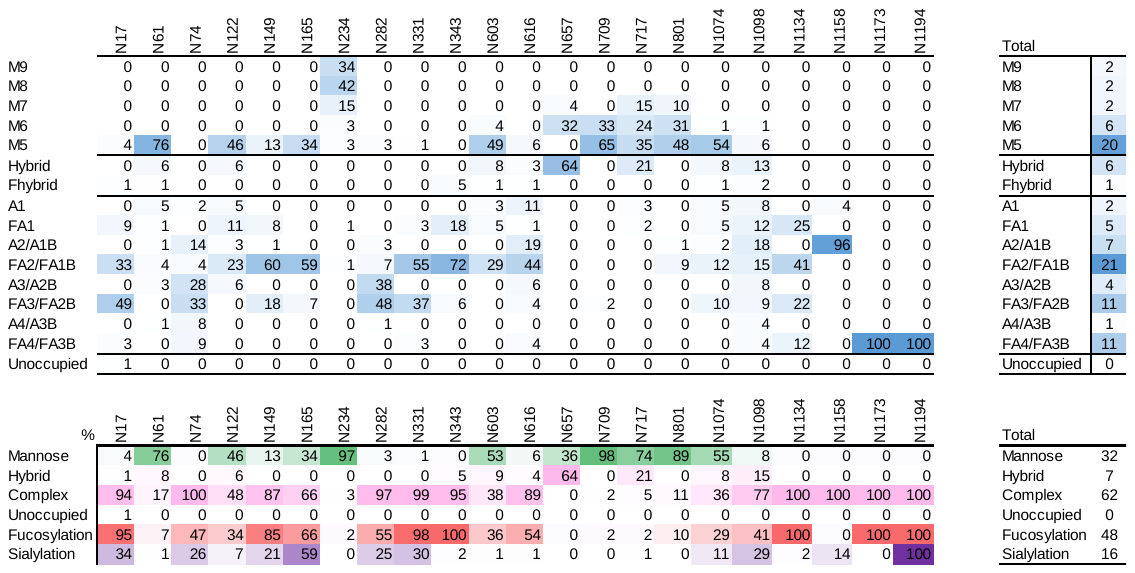

**Supplementary Table 1. Glycoform abundances observed across SARS CoV-2 S protein.** The upper table shows the categorized glycan compositions at each N-linked glycan site, with the global averages shown in the right-hand table. The lower table further categorizes the glycan compositions into oligomannose-, hybrid-, and complex-type
